# Trade-Offs between Speed, Accuracy, and Dissipation in tRNA^Ile^ Aminoacylation

**DOI:** 10.1101/2020.04.06.027615

**Authors:** Qiwei Yu, Joel D. Mallory, Anatoly B. Kolomeisky, Jiqiang Ling, Oleg A. Igoshin

## Abstract

Living systems maintain a high fidelity in information processing through kinetic proofreading, a mechanism to preferentially remove incorrect substrates at the cost of energy dissipation and slower speed. Proofreading mechanisms must balance their demand for higher speed, fewer errors, and lower dissipation, but it is unclear how rates of individual reaction steps are evolutionary tuned to balance these needs, especially when multiple proofreading mechanisms are present. Here, using a discrete-state stochastic model, we analyze the optimization strategies in *Escherichia coli* isoleucyl-tRNA synthetase. Surprisingly, this enzyme adopts an economic proofreading strategy and improves speed and dissipation as long as the error is tolerable. Through global parameter sampling, we reveal a fundamental dissipation-error relation that bounds the enzyme’s optimal performance and explains the importance of the post-transfer editing mechanism. The proximity of native system parameters to this bound demonstrates the importance of energy dissipation as an evolutionary force affecting fitness.

**Graphical TOC Entry:** 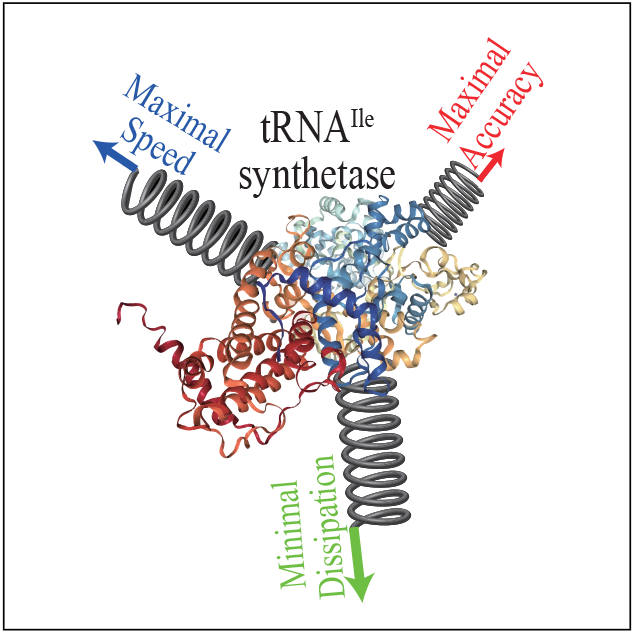

---

The fitness of many organisms relies on the fidelity and reliability of information propagation, epitomized by remarkable accuracy in DNA replication, mRNA transcription, and protein translation.^1–5^ It is well known that many biological systems adopt kinetic proofreading (KPR), a mechanism that actively removes wrongly incorporated substrates, to reduce the error beyond the equilibrium limit.^6,7^ The KPR mechanism is not only vital to cell viability,^8,9^ but also of great interest in the field of nonequilibrium thermodynamics.^10–13^

Ideally, biological systems should process information rapidly while keeping error and dissipation low. Theoretical studies, nonetheless, have revealed that it is impossible to optimize the three properties (speed, error, and dissipation) simultaneously.^14,15^ For example, copying of a single bit of information achieves its highest accuracy in either a slow and quasiadiabiatic regime or a fast and dissipative one, demonstrating a trade-off between speed and dissipation.^11^ Similar trade-offs involving the error also exist.^13^ Resolving these trade-offs requires biological systems to prioritize different properties. For replication and translation, enzymes were shown to optimize speed over error and dissipation.^14,16^ It is, however, unclear whether the same preference applies to more complex proofreading systems. For example, many enzymes exhibit multi-stage proofreading, i.e., the enzyme-substrate complex can undergo proofreading and be reset to the initial state at more than one state. Previous studies on multi-stage proofreading have focused on systems with discrimination between cognate and noncognate substrates only in dissociation steps.^10,12^ However, experimental evidence suggests that the discrimination can occur in any step.^4,5,17–20^ Thus, a systematic investigation of the speed-accuracy-dissipation relation in a biologically relevant context is still lacking.

Isoleucyl-tRNA synthetase (IleRS) in *Escherichia coli* is one of the best characterized multi-stage proofreading systems. During protein synthesis, IleRS must accurately pair cognate tRNA^Ile^ with the corresponding amino acid, i.e., isoleucine.^21^ Misincorporation has severe physiological impacts, including increasing the DNA mutation rate.^22^ IleRS utilizes multiple proofreading pathways to remove noncognate valine, an amino acid chemically similar to isoleucine. The proofreading mechanisms can be divided into pre-transfer editing, which occurs within the active (synthetic) site, and post-transfer editing, which occurs at a separate editing site. ^21,23^ In this work, we construct a biologically relevant model of IleRS (see Figure 1) and extract parameters from experimental data to address three questions: (i) how are the rates of individual reaction steps optimized to deal with the speed-error-dissipation trade-off? (ii) is there a fundamental limit on the extent of overall optimization, and if so, where is *E. coli* IleRS relative to that limit? (iii) is there an evolutionary reason for the existence of multiple proofreading pathways?

**Figure 1:**
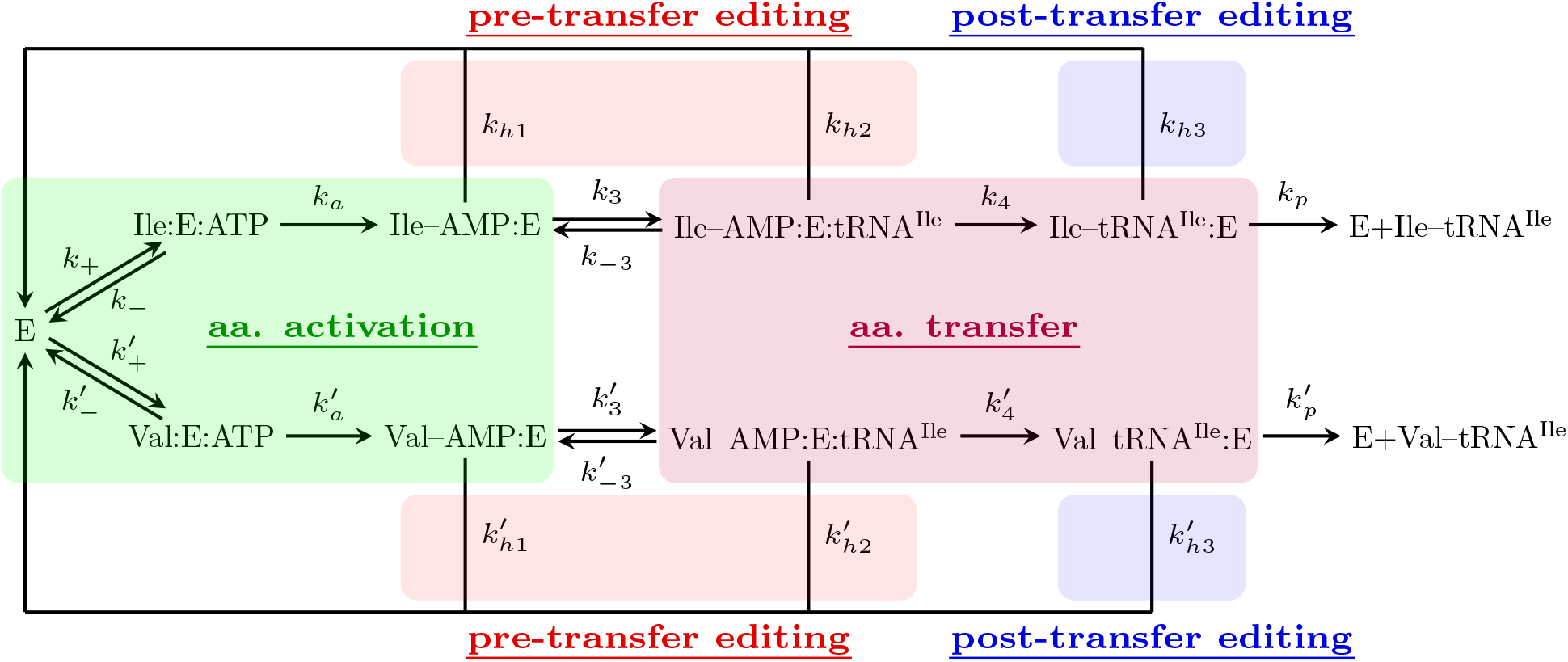
Chemical reaction network for the aminoacylation of tRNA^Ile^ in *E. coli*. Abbreviations: E, isoleucyl-tRNA synthetase (IleRS); Ile, isoleucine; Val, valine. The rate constants are labeled *k_i_* for the right pathway (isoleucine pathway) and 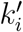 for the wrong pathway (valine pathway). The three proofreading pathways are labeled as *k*_*h*1_, *k*_*h*2_, *k*_*h*3_, and their primed counterparts. Although the reverse reactions are not drawn for some reactions which are kinetically driven forward, they are technically all reversible. We include them in the simulation to maintain thermodynamic consistency.

To this end, we quantitatively study the IleRS network using a discrete-state stochastic framework previously employed in the study of T7 DNA polymerase and *E. coli* ribosome.^14,16^ The chemical steps in Figure 1 are modeled as quasi-first-order transitions between different states with rates estimated from quantitative kinetic experiments^5,19,24–37^ (see SI for details). We define the error (*η*) as the ratio of the splitting probability of forming an incorrect product to the splitting probability of forming a correct product. The speed is quantified by inverse of the conditional mean first-passage time (MFPT, *τ*) to form a correct product starting from the free enzyme state (E). The dissipation is defined as the amount of free energy dissipated per product formed (*σ*, in units of *k_B_T*).^38^ The detailed mathematical definitions and procedures can be found in the SI.

Due to the symmetry of the reaction network, each process for the noncognate amino acid (with rate 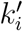) has a corresponding reaction in the cognate reaction network (with rate *k_i_*). We relate those two reactions by defining a set of discrimination factors 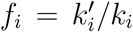, where *i* is a subscript unique to individual reactions. The discrimination factors are the fundamental reason that differentiates the cognate reactant from the non-cognate one. We study the interplay between three characteristic properties (speed, error, and dissipation) by proportionally varying the rate constants (*k_i_* and 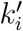) while keeping the discrimination factors *f_i_* fixed. In this way, neither of the substrates receives any unfair advantage. A trade-off between two properties occurs if a change in a given rate constant cannot simultaneously improve both of them. Thus, the trade-offs between speed, error, and dissipation must be discussed in the context a specific set of rate constant perturbations. We investigate the effect of either tuning one rate constant (local parameter perturbation) or tuning multiple rate constants (global parameter perturbation).

We begin our analysis with two catalytic steps: amino acid activation and transfer, denoted by *k_a_* and *k*_4_, respectively (see Figure 1). These two processes are energetically important since they involve covalent-bond breakage and formation, thereby channeling the energy stored in the phosphoanhydride bonds of ATP to that in the ester bond of aa-tRNA. The energy transferred provides the driving force for subsequent peptide bond synthesis in the ribosome. To uncover the optimization for these steps, we analyze how the characteristic properties change as a function of their respective rate constants (*k_a_* and *k*_4_)

The results shown in Figure 2 are somewhat unexpected. First, both speed and accuracy can be improved simultaneously by increasing the catalytic rates of both reactions: this result demonstrates a lack of trade-off between the two properties (Figure 2a, c).Indeed, the synthetase has a higher affinity for isoleucine over valine, which results in slightly higher catalytic rates compared to that of valine. In other words, the discrimination factors *f_a_* and *f*_4_ are slightly smaller than 1. ^19^ Therefore, speeding up these steps magnifies the discrimination in these two steps and results in higher speed and accuracy for IleRS. Second, the relation between speed and dissipation is non-monotonic (Figure 2b, d). Consequently, the MFPT–dissipation curve can be separated into two branches: a trade-off branch where reducing dissipation inevitably increases MFPT, and a non-trade-off branch where the two properties can be improved simultaneously. However, for both steps, the native system (i.e., the green dot corresponding to experimentally measured rates of these steps) resides on the tradeoff branch. The fact that the native system also resides between the minima of error and dissipation (magenta and pink squares) indicates a trade-off between these two properties as well. Therefore, it is the dissipation that prevents the co-optimization of speed and accuracy. This result is analogous to prior findings for the Pol-Exo sliding in T7 DNA polymerase and the proofreading rate in aa-tRNA selection by *E. coli* ribosome.^14^

**Figure 2:**
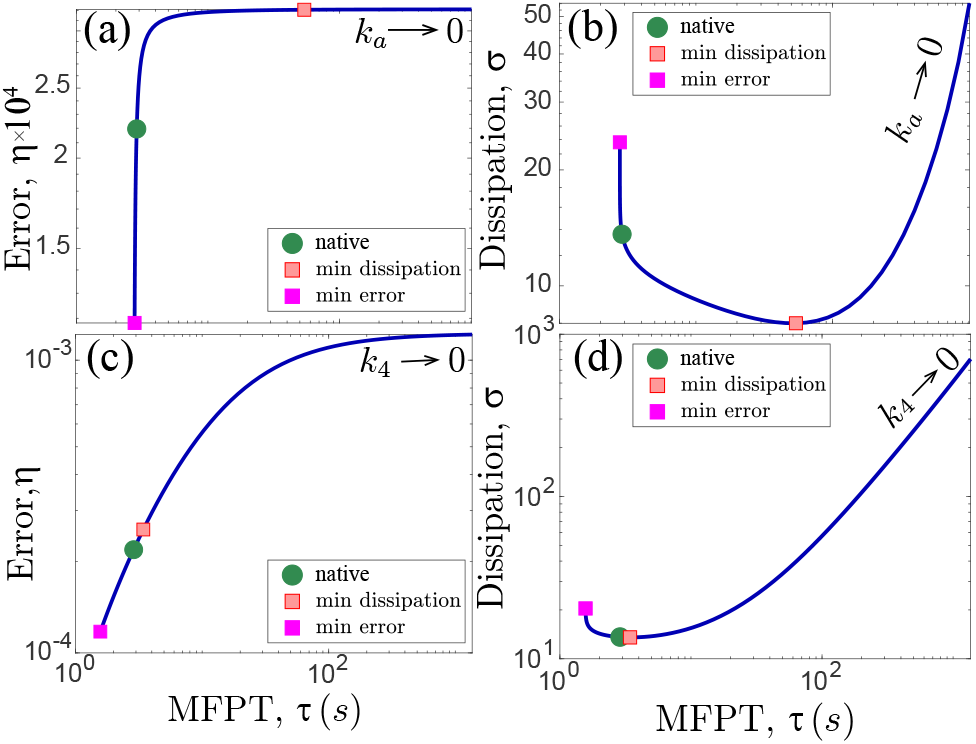
The trade-off between MFPT (*τ*), error (*η*), and dissipation (*σ*) due to variation of the two key catalytic steps: amino acid activation coupled by ATP hydrolysis (*k_a_*, upper panels), amino acid transfer (*k*_4_, lower panels). Catalytic rates for the non-cognate substrates (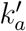 and 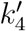) are varied proportionally to keep the discrimination factors fixed. The green dot denotes the native system. The pink and magenta squares correspond to the positions of the minimum dissipation and error, respectively. The termination of the curve at the magenta squares is not a result of insufficient sampling range, but a limit which cannot be passed even when the rates go to infinity.

Qualitatively, the trade-off mainly occurs between speed and dissipation. Due to the low specificity in activation and transfer,^19^ the change in error due to the variation of these parameters (Figure 2a, c) is actually marginal, especially compared to its variation when the the downstream quality control steps are varied (Figure 3b, d). Nevertheless, the two reactions take strikingly different strategies in dealing with the trade-off. The activation step prefers to optimize the speed, while the transfer step is near minimal dissipation. The activation step is not rate-limiting, i.e., the maximal speed achieved at *k_a_* → ∞ is only 4% larger than the native value. However, such increase in speed leads to a drastic (70%) decrease in dissipation, as evident from a nearly vertical slope of the curve connecting green dot and magenta square in Figure 2b. On the other hand, further decreasing the rate of this step would make it rate-limiting as reducing the dissipation by 41% will increase the MFPT by 18 fold. We hypothesize that the evolution tunes the activation reaction to be on the cusp of being rate-limiting, i.e., to decrease dissipation while keeping the impact on the speed marginal.

**Figure 3:**
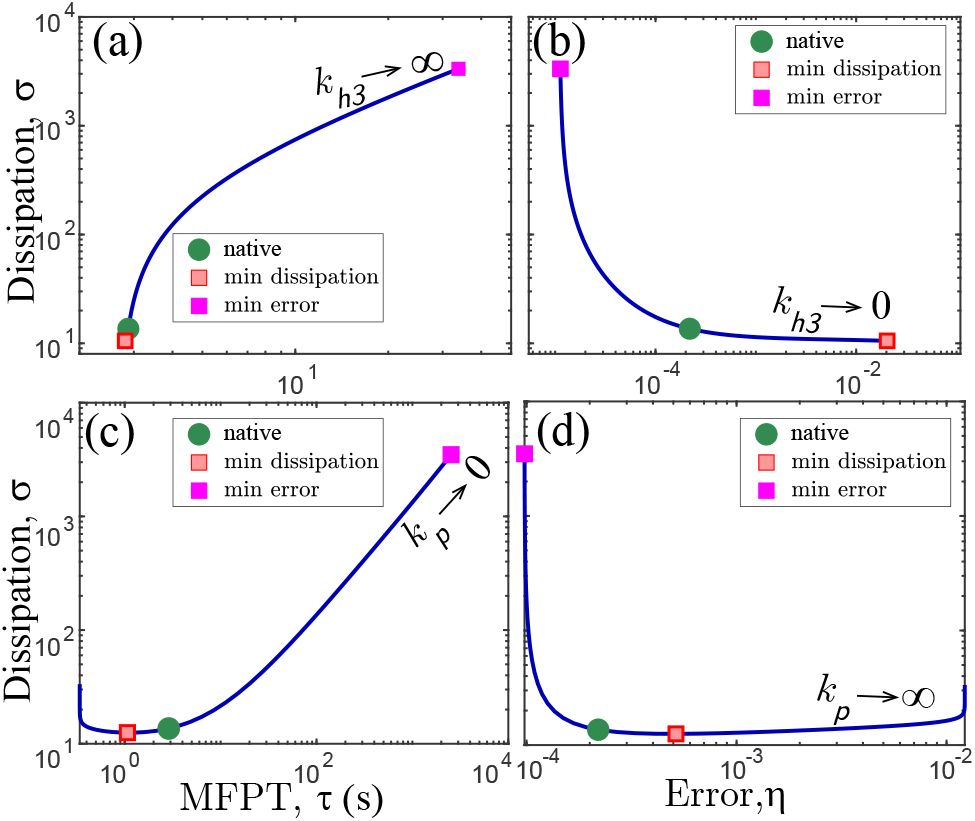
The trade-off between MFPT (*τ*), error (*η*), and dissipation (*σ*) due to variation of the two key quality control processes: the translocation and deacylation of aa-tRNA (*k*_*h*3_, upper panels) and product release (*k_p_*, lower panels). Catalytic rates for the non-cognate substrates (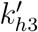 and 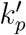) are varied proportionally to keep discrimination factors fixed. The green dot denotes the native system. The pink and magenta squares correspond to the positions of the minimum dissipation and error, respectively. When *k_p_* → 0 or *k*_*h*3_ → ∞, both the MFPT and dissipation diverge to infinity, while the error rate converges to a constant.

In stark contrast, the transfer step rate *k*_4_ is nearly optimized for minimal dissipation—the green dot in Figure 2d is only 0.5% above the minimal value denoted by the pink square. An increase in the transfer rate can reduce the MFPT by 45%, but this will lead to a 50% increase in the dissipation. Thus, the increase of MFPT due to optimizing the dissipation in the transfer step is considerably smaller as compared to 18-fold MFPT increase in the activation step (Figure 2b). This finding makes optimization of dissipation in the transfer step more plausible.

Altogether, the results on Figure 2 demonstrate distinct evolutionary optimization strategies in two catalytic steps. IleRS tunes up the amino acid activation rate to guarantee fast production of aa-tRNA and keeps the transfer rate at an intermediate level so that the dissipation is nearly minimal. These strategies employed enable partial reconciliation between the speed-dissipation trade-off for the enzyme and allow for faster and more efficient formation of aa-tRNA.

Next, we focused on the optimization strategies in key quality control steps. The ability of IleRS to preferentially hydrolyze misacylated products is largely determined by the rate constants for post-transfer editing (*k*_*h*3_) and product release (*k_p_*). The two pre-transfer editing steps exhibit behaviors similar to post-transfer editing, but they are less effective in the ability to suppress the error.^20^ Following the same methodology, we present the results in Figure 3. In contrast to those of activation and transfer, the native rates of the quality-control steps reside on a non-trade-off branch of the speed-dissipation curve. For these steps, it is the error that prevents the co-optimization of speed and dissipation. Indeed, improving the accuracy requires either decreasing *k_p_* or increasing *k*_*h*3_ and hence, increasing the proofreading fluxes. Consequently, more ATP is consumed without forming aa-tRNA and more time will be needed to successfully form a product.

We further explore how the system balances the need for higher accuracy and the resultant cost in terms of slower speed and higher dissipation by examining the position of the native system on the trade-off plots. We found that, in both cases, the native system resides not far away from the min-dissipation/min-MFPT point. For example, the MFPT of the native system is only 2% above the minimum due to variation of *k*_*h*3_, and the dissipation in the *k_p_* trade-off plot is only 9% above the minimum value. Although it is hard to claim that either speed or dissipation is the “deal-breaker” which renders further improvement of the accuracy unfavorable in evolution, it seems that their combined effect results in a compound cost that decreases the organism’s overall fitness. Consequently, both *k*_*h*3_ and *k_p_* are tuned in a way that the MFPT and dissipation are optimized as much as possible as long as the error rate is within a reasonable range, i.e., ~ 10^−4^. Indeed, the fidelity of the aminoacylation only needs to surpass the overall accuracy of protein synthesis (*η* = 10^−3^ − 10^−4^).^3–5^ Further improvements beyond this threshold will not significantly suppress the protein synthesis error since more mistakes will be made during aa-tRNA selection at the ribosome. Moreover, the accuracy can be further improved through downstream error-correcting mechanisms such as EF-Tu specificity. The need to synthesize numerous proteins continuously raises the necessity of decreasing the cost of time and energy to deliver a single amino acid while keeping the error rate tolerable. Collectively, these factors rationalize our finding that the proofreading mechanism in *E. coli* IleRS synthetase has evolved to adopt an “economic” error correcting strategy which establishes a reasonable level of fidelity in a speedy and inexpensive fashion.

Our analysis of individual reactions has shown that although different steps can adopt different optimization strategies, with most reaction steps optimize IleRS towards an energetically efficiency (i.e., minimal dissipation). However, both our data and previous estimations in the literature suggest that over 10% of the ATP hydrolyzed leads to proofreading fluxes, which do not form any products.^39^ It is therefore unclear whether these futile fluxes are inevitable and whether the local optimization strategies collectively result in the global optimization of dissipation. To address this question, we perform global parameter sampling. To this end, all of the kinetic parameters are varied under the constraints of fixed discrimination factors and fixed chemical potential differences for both futile and product formation cycles (i.e., Δ*μ*_ATP_ and Δ*μ_p_*, respectively). The sampling can be regarded as a generalization of the local trade-off analysis. As shown in Figure 4, we found that all sampled systems always reside on one side of a boundary (blue dashed line) on the error-dissipation plane. Thus, this boundary is a fundamental constraint imposed by the discrimination factors which cannot be circumvented through any variation of the kinetic parameters. The minimum amount of dissipation for any level of accuracy is Δ*μ_p_* = 9.8 *k_B_T*, which corresponds to the absence of any proofreading. Towards the other end of the spectrum, the error rate can be suppressed to as low as *η*_min_ ~ 6 × 10^−7^, but the dissipation increases as the error rate decreases and diverges to infinity as the error approaches its minimum. Our results reaffirm the notion that an increased amount of free energy must be dissipated to enhance biological fidelity.^6,15^ On the other hand, the sampling does not put any constraint on the optimal speed, since the timescale of the system can be tuned arbitrarily by varying all the rate constants proportionally. In reality, however, the kinetic rates are limited by the underlying biochemical processes. For instance, it is hard to conceptualize how the binding rates of ATP and amino acid to IleRS can exceed the diffusion limit.^40^ These biochemical constraints could reduce the parameter space reachable for the actual system, resulting in a sub-optimal performance compared to the theoretical bound discussed here.

**Figure 4:**
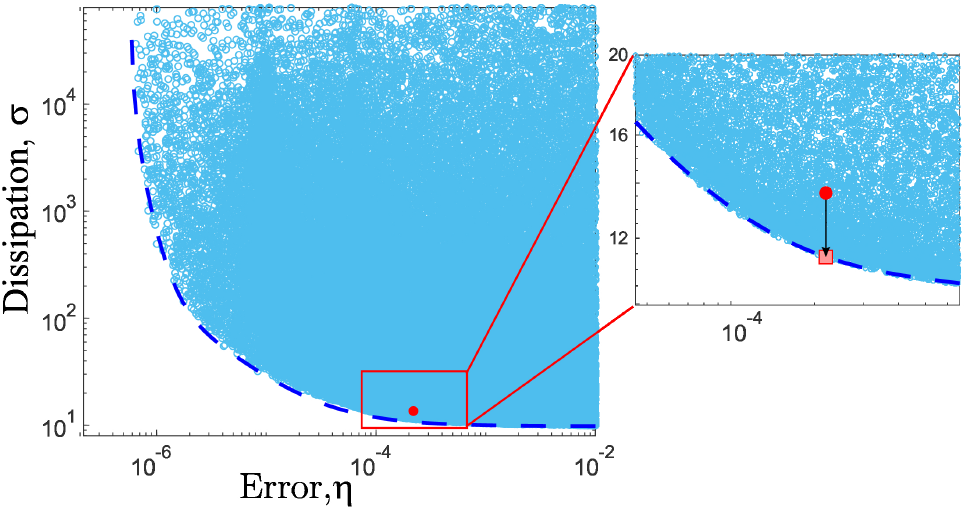
The scatter plot of the error-dissipation relation resulting from the variation of all rate constants. The area to the left or under the blue dashed boundary is inaccessible under any combinations of the rate constants. The red dot indicates the native system; the pink square marks the theoretical minimum dissipation for the native accuracy.

The existence of a global lower bound on dissipation for any given error underscores the nonequilibrium nature of biological information processing. In this system, active proofreading resets the system into the initial state without forming a product and therefore, results in futile ATP hydrolysis. Such proofread fluxes can only improve the system’s accuracy if the futile cycles are powered by free energy released from ATP hydrolysis. Thus, an increase in proofreading frequency inevitably leads to an increase in dissipation. This trade-off is evidenced by the global dissipation bound that decreases with error (see Figure 4). A similar dissipation-accuracy relation was also found in a system for sensory adaptation, where the optimal dissipation increases with adaptation accuracy and eventually diverges at perfect adaptation, a behavior similar to the bound found here. ^41^ In both cases, dissipation is associated with cyclic futile fluxes that consume free energy to drive the system away from thermodynamic equilibrium. In contrast, the proofreading fluxes in an equilibrium proofreading system (i.e. without any free energy yield from ATP hydrolysis) will not reduce the error of the system.

A close-up inspection at the lower right area of Figure 4 reveals that the native system resides in close proximity to the error-dissipation boundary. Indeed, the dissipation can at most be decreased by only 16% (~ 2.2 *k_B_T*) without any reduction in accuracy (i.e., from the red dot to the pink square on the Figure 4 inset). As compared to the complete dynamic range of error and dissipation, this is a very small margin. According to the error-dissipation bound, the dissipation must be increased by three-fold (to 34 *k_B_T*) in order to reduce the error rate by one order of magnitude (from 10^−4^ to 10^−5^). This increase is a tremendous cost considering the amount of amino acid (in this case, isoleucine) required for protein synthesis in the *E.coli* cell cycle. It is therefore understandable that the synthetase has chosen an economic strategy in quality control, as seen in the trade-off analysis of *k*_*h*3_ and *k_p_*, instead of promoting its accuracy to an unnecessary level.

To understand the evolutionary necessity of multi-stage proofreading in *E. coli* IleRS, we use our model to assess the importance of different proofreading mechanisms. As shown in Figure 5a, removing both pre-transfer editing pathways causes a marginal increase in error (5%, from purple bars to yellow bars), while removing the post-transfer editing pathway increases the error drastically (by two orders of magnitude). This result is consistent with experimental findings that post-transfer editing contributes to the majority of the editing in *E. coli* IleRS.^5^ The pre-transfer proofreading fluxes takes up only 12% of all editing fluxes. Notably, this estimate is smaller than the 30% derived from a previous kinetic experiment that employed the Michaelis-Menten equation to fit the AMP formation rate.^20^ This discrepancy is likely because our framework takes into account a more realistic mechanism of the reaction (as compared to Michaelis-Menten).

**Figure 5:**
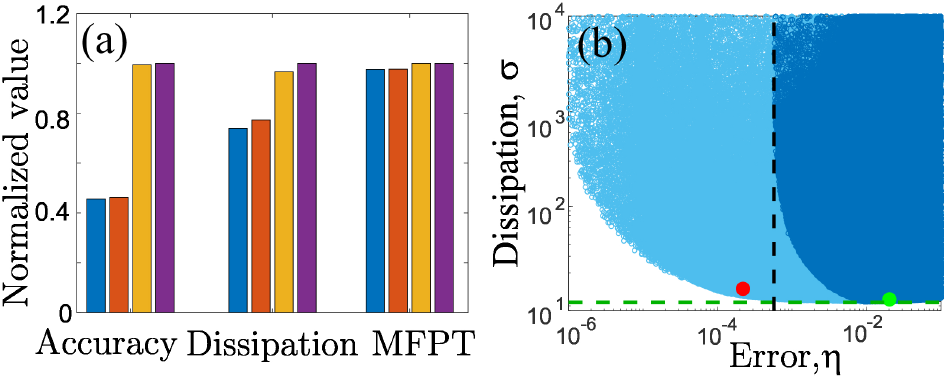
Quantitative comparison of the importance of different proofreading pathways. Left: Key properties for four variants of IleRS. Blue, no proofreading; orange, pre-transfer editing only; yellow, post-transfer editing only; purple, the native system. The accuracy is defined as − ln *η*. All values are normalized by taking the ratio against the native system. Right: error-dissipation scatter plot due to global parameter variation. Dark blue, samples without post-transfer editing; light blue, samples with wild-type network; red dot, the native system (*η*_nat_ = 2.2 × 10^−4^); green dot, mutant with only pre-transfer editing (*η*_natp_ = 2.0 × 10^−2^). The black dashed line indicates the minimum error for the system without post-transfer editing (*η*_min_ = 5.5 × 10^−4^). The green dashed line stands for the minimum dissipation (*σ*_min_ = *μ_p_* = 9.8 *k_B_T*)

Given the importance of post-transfer editing, it is intriguing to explore the specific need that caused the evolution of this mechanism. To investigate this, we performed global parameter sampling in the absence of post-transfer editing 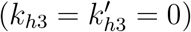 and superimposed the error-dissipation relation onto the native one (see Figure 5b). The addition of post-transfer editing significantly expands the reachable area on the error-dissipation plane. It not only allows IleRS to achieve a higher accuracy, but also makes it energetically more efficient to maintain an accuracy level attainable by pre-transfer editing alone. For each of the optimized systems on the boundary of the dark blue area, there is a system in the light blue area that can achieve the same accuracy with a much lower dissipation. Somewhat surprisingly, the results indicate that even without post-transfer editing, the minimal error (*η*_min_ = 5.5 × 10^−4^, black dashed line in Figure 5b) is just about 2.5-fold higher than in the native system (*η*_nat_ = 2.2 × 10^−4^, red dot in Figure 5b). As discussed above, this level of accuracy could be sufficient. However, the amount of dissipation corresponding to this error rate is astronomical. For example, to reach the error rate 3-fold higher than that of the native system (3*η*_nat_ = 6.6 × 10^−4^), the dissipation must be increased to at least 200 *k_B_T*. Further decrease of the error requires more and eventually an infinite amount of dissipation as shown by the asymptotic behavior of the boundary of the dark blue area.

Therefore, our results here provide a quantitative rationalization of the evolutionary origin of the IleRS CP1 editing domain. The low discrimination factor in pre-transfer editing (*f*_*h*1_ and *f*_*h*2_) makes it energetically costly to maintain a higher accuracy as shown by the accessible region (the dark blue area) in Figure 5b. Since both amino acid activation and pre-transfer editing take place within the active site, it might be improbable to improve the pre-transfer editing specificity any further without affecting the transfer efficacy. Consequently, a separate domain is recruited for post-transfer editing, which provides a more economic way of maintaining genetic code fidelity. The existence of multiple proofreading pathways successfully improves the overall fitness by relaxing the trade-off between error and dissipation. Consistent with this prediction, a high degree of conservation in CP1 domains for IleRS, ValRS, and LeuRS proteins suggests early emergence and selective pressure to maintain post-transfer editing.^42^

To summarize, our biophysical model with parameters derived from experiments leads to valuable insights on the interplay between speed, accuracy, and dissipation for the isoleucyl-tRNA synthetase. As a prerequisite to peptide synthesis at the ribosome, it is essential to maintain a high speed and energetic efficiency of aa-tRNA synthesis. However, we discovered that their co-optimization is prevented by a dissipation-speed trade-off in key catalytic steps. Generalizing the previous viewpoint on the existence of a universal preference to improving some properties at the cost of the others,^16^ we provide a new perspective that different priorities of individual reaction steps enable partial reconciliation of the trade-offs.

Despite its potential to reduce the error significantly (by two orders of magnitude, see Figure 4), it seems that the IleRS has adopted an “economic” strategy which improves speed and dissipation as much as possible and maintains the error at a reasonable level. We demonstrate the extent to which dissipation is optimized by the proximity of the native system to a fundamental error-dissipation bound. The results are further rationalized by several biological arguments, including the error threshold imposed by the downstream aa-tRNA selection (*η* = 10^−3^ − 10^−4^) and the demand for rapid and energetically inexpensive aa-tRNA supply for protein synthesis. Once the accuracy of aaRS surpassed that of aa-tRNA selection, evolutionary forces no longer drove any enhancement of accuracy. Instead, reducing MFPT and dissipation became the priorities for IleRS. This hypothesis could be experimentally tested by investigating whether the dissipation can be reduced in IleRS mutants without impairing the overall fidelity of protein translation.

Our results not only provide new understandings of the aminoacylation process, but also construct a general framework for analyzing complicated discriminatory proofreading networks. We argue that the importance of one reaction pathway can never be fully appreciated by looking at one property in isolation. Instead, one should examine the accessible region in the space spanned by the characteristic properties such as the error-dissipation plane studied here. From this perspective, the significance of post-transfer editing lies not in reducing the minimum error rate, but in making it less energetically costly to maintain an error rate of *η* ~ 10^−4^. The existence of a dissipation-error lower bound (blue-dashed line, Figure 4) indicates a minimum cost for biological error correction that is imposed by the discrimination factors. Further studies on this minimum energy cost may provide new insights to the underlying thermodynamic principle of biological information processing.

## Acknowledgments

This work was supported by Welch Foundation Grant Award C-1995 to O.A.I. and Center for Theoretical Biological Physics National Science Foundation (NSF) Grant PHY-1427654. A.B.K. also acknowledges support from the Welch Foundation Grant C-1559. Q. Y. acknowledges support from Chinese Scholarship Council (CSC).

## Associated Content

### Supporting Information

Detailed description of the tRNA^Ile^ aminoacylation model (including the methods for estimation of the kinetic parameters); mathematical formalism and definitions of the error rate *η*, MFPT *τ*, and the energy dissipation *σ*.

## Supporting Information

### I. CHEMICAL NETWORK FOR THE tRNA^Ile^ AMINOACYLATION

In this section, we provide a more detailed account on our model of the mechanism of aminoacylation by IleRS, including the derivation of the biochemical parameters in the model. The aminoacylation mechanism can be divided into two stages [1, 2]. In the first stage, the amino acid (aa) is activated at the active site of the synthetase by hydrolyzing ATP to form aminoacyl-adenylates (Ile-AMP or Val-AMP), as shown by the reactions in the green box (see Figure S1). The tRNA can subsequently bind to the complex. The charged amino acid is then transferred to the 3’ end of the tRNA, which is represented by the reactions in the purple box (see Figure S1). Finally, the charged aminoacyl-tRNA dissociates from the synthetase at rate *k_p_* (or 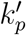) and is delivered to the ribosome by the elongation factor Tu (EF-Tu). Since we are mainly concerned with the accuracy of tRNA charging, the delivery and any subsequent process will not be included in the model.

The amino acid substrate specificity is achieved not only by the preferential binding of the cognate amino acid, but also through selective editing mechanisms [2, 3]. Although these proofreading pathways mainly serve to reduce the production rate of misacylated tRNA, they can also occur (albeit with lower probability) for complexes with the right amino acid. Therefore, we construct a completely symmetric reaction network where all proofreading mechanisms exist for both amino acids. The preference for the cognate substrate is reflected in the higher affinity and lower editing rates. Specifically, there are two pre-transfer editing pathways (red boxes in Figure S1) and one post-transfer editing pathway (blue boxes in Figure S1). The first pre-transfer editing (modeled by *k*_*h*1_) is tRNA-independent and occurs before tRNA binding. In this reaction, the aminoacyl-adenylate (aa-AMP) is either dissociated from the synthetase or hydrolyzed at the active site. We do not distinguish dissociation from hydrolysis because the synthetase is reset to its initial state (E) after either reaction. The second pre-transfer editing pathway (modeled by *k*_*h*2_) is tRNA-dependent and hydrolyzes the aa-AMP complex before the amino acid could be transferred to tRNA, which is already bound to the complex. Finally, the post-transfer editing pathway (modeled by *k*_*h*3_) occurs after the transfer and results in the translocation of aminoacyl-tRNA from the active site to the editing site and its subsequent hydrolysis (deacytlation). We are only concerned with the proofreading of aminoacyl-tRNA that occurs prior to its release from the aminoacyl-tRNA synthetase. Quality control mechanisms that take place after the aminoacyl-tRNA is released (e.g., the preferential binding of the elongation factor Tu to the correctly acylated-tRNA[2]) constitute a different layer of specificity enhancement and hence, are not explicitly considered in the model. The model focuses on the stochastic dynamics of a single isoleucyl-tRNA synthetase and assumes constant concentrations of other molecular species in the cellular environment such as amino acids. Although the low activation energy for the hydrolysis of the ester bond of aa-tRNA allows for the deacylation of misacylated products by either editing site residues or the free-hydroxyl groups of A76 [2], they are not relevant in this current framework.

**FIG. S1.**
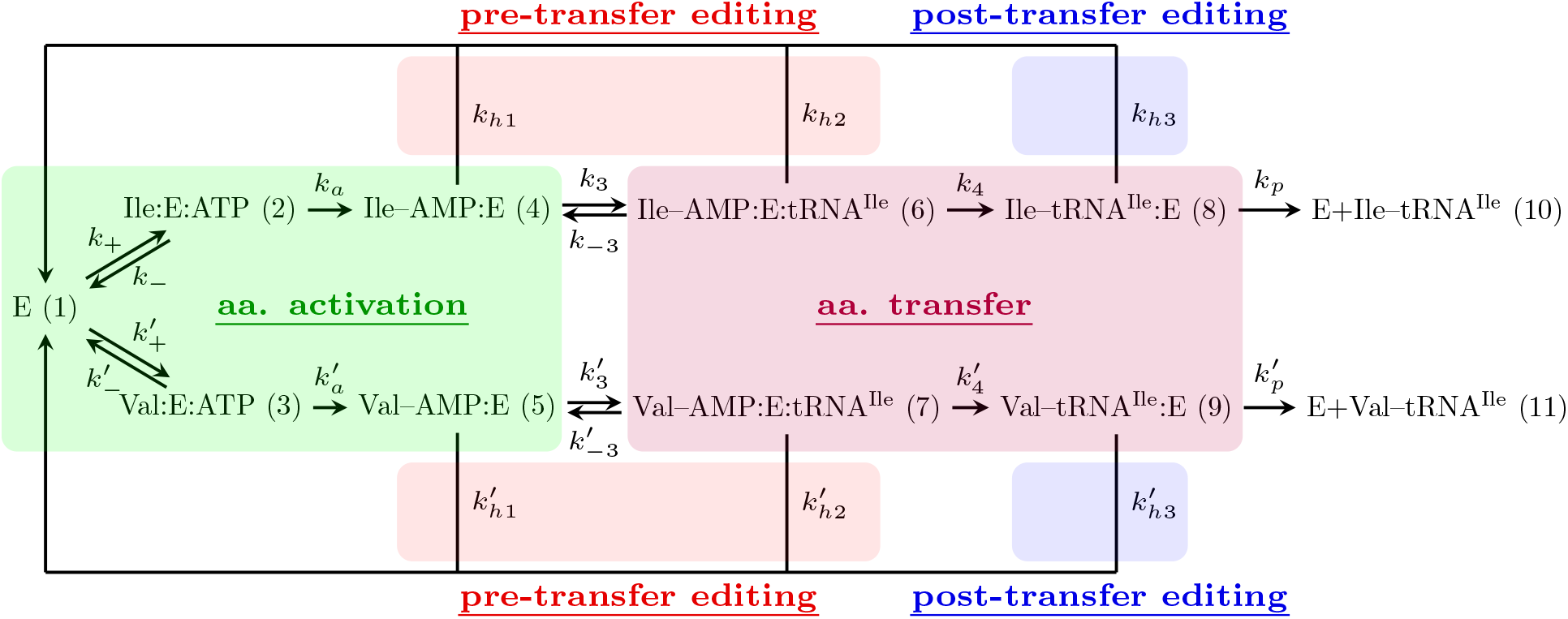
Chemical reaction network for the aminoacylation of tRNA^Ile^ in *E. coli*. Abbreviations: E, isoleucyl-tRNA synthetase (IleRS); Ile, isoleucine; Val, valine. The rate constants are labeled *k_i_* for the right pathway (isoleucine pathway) and 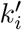 for the wrong pathway (valine pathway). The three proofreading pathways are labeled as *k*_*h*1_, *k*_*h*2_, *k*_*h*3_, and their primed counterparts. The chemical states are numbered from 1 to 11 as indicated by the numbers in the parentheses.

The above-described mechanism is illustrated on Figure S1 and serves as foundation for our biophysical model. The rate parameters are either directly obtained or indirectly derived from published kinetic experiments [3–18] as described below.

#### A. Determination of Rate Constants from the Literature

The numeric values and sources of all rate constants that are shown in Figure S1 are summarized in Table S1. Most of them are directly obtained from the literature. Explanations for a few of parameters that are indirectly derived or estimated are listed below:

- The tRNA binding steps (*k*_3_ and 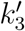) are assumed to be diffusion-limited. Their forward rates are estimated by the Smoluchowski equation *k* = 4*πDR*, where *D* and *R* are the diffusion coefficient and the radius of the molecule. There backward rate constants are calculated as the product of the dissociation constant and the forward rate constant.
- The binding of amino acid and ATP are model as one step with the assumption that ATP binding is diffusion-limited. The detailed procedures are given in the next subsection.
- Product release rate is derived by using the the three slowest steps to estimate the waiting time 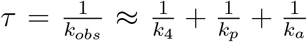, from which *k_p_* can be solved. *k_obs_* is given by Table 4 of ref. [13].
- The product release rate for the non-cognate substrate 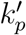 is assumed to be the same as the cognate product release rate *k_p_* based on a chemical argument. Given that valine and isoleucine differ only in the side chain (which is a small part of the whole aminoacyl-tRNA complex) and that the amino acid has been transferred to the tRNA, we argue that this difference should not affect the product release rate.

**TABLE S1.** Estimated first-order kinetic parameters involved in the tRNA^Ile^ aminoacylation model.

| Parameter | Value ($\text{s}^{-1}$) | Sources |
| --- | --- | --- |
| $k_+$ | $6.2 \times 10^5$ | [3, 5, 14, 15] |
| $k_-$ | 30 | [3, 5, 14, 15] |
| $k'_+$ | $7.0 \times 10^3$ | [3, 5, 14, 15] |
| $k'_-$ | 28 | [3, 5, 14, 15] |
| $k_a$ | 40 | $k_{cat}$ from [3] |
| $k'_a$ | 31 | $k_{cat}$ from [3] |
| $k_3$ | $8.7 \times 10^2$ | [8, 17, 18] |
| $k_{-3}$ | $1.1 \times 10^2$ | [4, 17, 18] |
| $k'_3$ | $8.7 \times 10^2$ | [8, 17, 18] |
| $k'_{-3}$ | $1.1 \times 10^2$ | [4, 17, 18] |
| $k_4$ | 3.8 | Figure 2 of [3] |
| $k'_4$ | 2.3 | Figure 2 of [3] |
| $k_{h_1}$ | 0.003 | Table 1 of [3]; Supp. Table 1 of [13] |
| $k'_{h_1}$ | 0.036 | Table 1 of [3]; Supp. Table 1 of [13] |
| $k_{h_2}$ | 0.0135 | Table 1 of [3] |
| $k'_{h_2}$ | 0.128 | Table 1 of [3] |
| $k_{h_3}$ | 0.058 | Table 4 of [3] |
| $k'_{h_3}$ | 64 | Table 4 of [3] |
| $k_p$ | 0.65 | Table 4 of [13] |
| $k'_p$ | 0.65 | Table 4 of [13] |

#### B. Justification for Modeling Amino Acid Binding and ATP Binding as One Step

The first step of our kinetic model (Figure 1) is the charging of the amino acid. It includes binding of both the amino acid and ATP to the enzyme followed by transfer of AMP to the amino acid and release of the pyrophosphate PP_i_. Generally speaking, the amino acid and ATP don’t have to bind sequentially. Here, we construct a one-step binding model with effective first-order reaction rates *k*_+_ and *k*_−_ under physiological conditions by exploring the complete model in which ATP or the amino acid bind in random order.

First, we consider a sequential binding model (*s*_1_ is the amino acid and *s*_2_ is ATP),

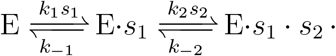

Using *x*, *y*, and *z* to denote the concentration of the three states of the enzyme, the quasisteady-state approximation of E·*s*_1_ reads:

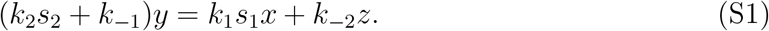

The net flux through this pathway is

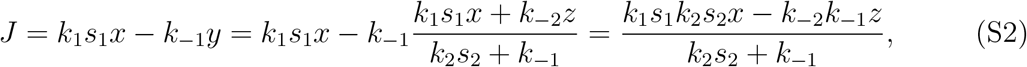

so we can view this model as a single step reaction with:

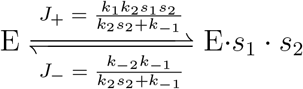

Note that the forward reaction rate is proportional to *s*_1_*s*_2_, so we can effectively write it as the simultaneous binding of *s*_1_ and *s*_2_. Since the *s*_1_ and *s*_2_ are completely symmetric in this case, we can write the other pathway in which *s*_2_ binds first in the same form and sum the reaction constants together. In the end, we have

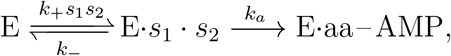

in which 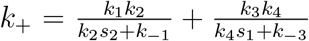 and 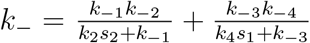 are both dependent on the ATP and amino acid concentrations. Mathematically, the total flux can be written in the Michaelis-Menten form

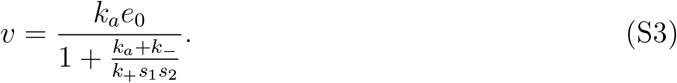

To get parameter estimations for this steps from experiments, we refer to Table 6 of ref. [3] where the production of the aa·E·AMP complex is measured by pyrophosphate exchange experiments. The authors fitted to a Michaelis-Menten scheme to their kinetics data with a standard method of determining the Michaelis-Menten constant through a double reciprocal plot (*k*_*a*_*e*_0_*v*^−1^ and 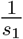 are plotted against each other). For our scheme, we have

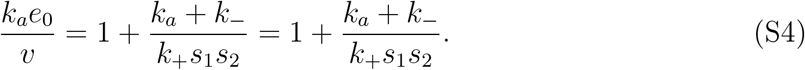

Note that *k*_+_ and *k*_−_ are both functions of *s*_1_ and *s*_2_, so the double reciprocal plot is not exactly a straight line. Since the concentration *s*_1_ was varied around *K_m_* in the experiment, the measured Michaelis-Menten constant should be the slope of the tangent near *s*_1_ = *K_m_*, mathematically written as:

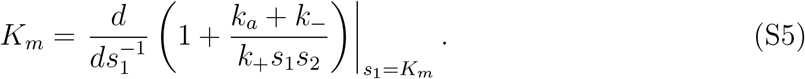

Effectively, we are doing a Taylor expansion of Equation S4 near *s*_1_ = *K_m_*, whose solution is

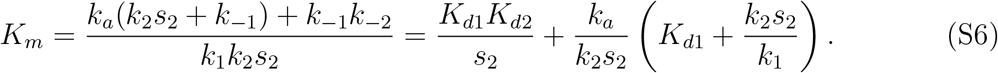

Assuming that ATP (*s*_2_) binding is diffusion limiting, the binding rate is

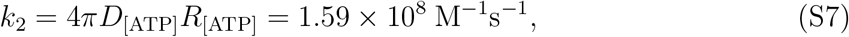

where *D*_ATP_ is obtained from [14]; *R*_ATP_ is obtained from ref. [15]. Considering the physiological concentration of ATP, we have *k*_2_*s*_2_ = 1.53×10^6^ s^−1^ and *k*_−2_ = *K*_*d*,ATP_*k*_2_ = 2.78×10^4^ s^−1^. On the other hand, the experiment measurements are *k*_cat_ = 40.2 s^−1^ and *K*_M_ = 6.9 *μ*M for isoleucine, and 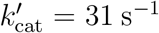 and 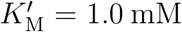 for valine. The concentration of ATP used was 4.0 mM. *k*_1_ can be explicitly solved from Equation S6:

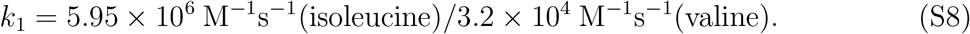

Hence, *k*_1_*s*_1_ = 1.8 × 10^4^ s^−1^(isoleucine)/1.3 × 10^2^ s^−1^(valine). The reverse rate constant *k*_−1_ = *k*_1_*K*_*d*1_ = 48 s^−1^(isoleucine)/28 s^−1^(valine).

Assuming symmetric rate constants, namely, *k*_±1_ = *k*_±4_ and *k*_±2_ = *k*_±3_, we then calculate the value of *k*_+_ and *k*_−_ under physiological concentrations and use this single step to represent the binding and unbinding of both substrates. The pseudo-first order rate constants are:

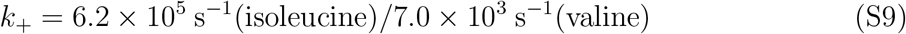

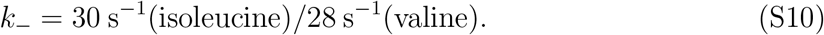

#### C. Thermodynamic Constraints and the Determination of Reverse Reaction Rates

Although some of the steps are strongly driven forward, we make all reactions reversible in the simulation for the sake of thermodynamic consistency as irreversible reactions always lead to infinite dissipation. The rate constants for reverse reactions are determined by a set of thermodynamic constraints that connect the ratio of the rate constants with the chemical potential difference of the reactants and products of any reaction cycle [19]:

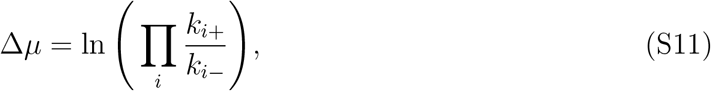

where *k*_*i*+_ and *k*_*i*−_ stands for the forward and backward rate constants; Δ*μ* stands for the chemical potential difference between the reactants and products. For futile cycles Δ*μ*_ATP_ = 29.5 *k_B_T*, and for product formation cycles Δ*μ_p_* = 9.8 *k_B_T*. In this system, the thermodynamic constraints read

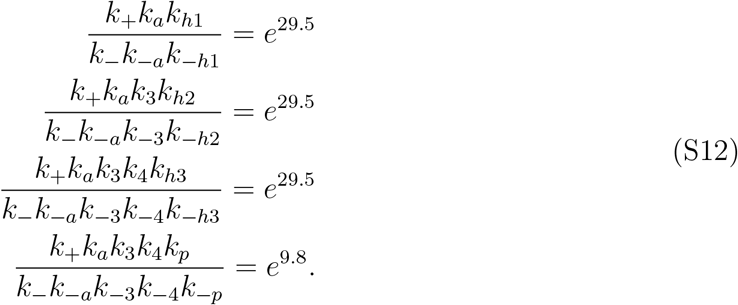

The same set of constraints also exist for the noncognate reactions, where the rate constants are simply replaced by their primed counterparts. The reverse rate constants that are omitted in Figure S1 can be determined from these relations. Their values are introduced purely for the calculation of the dissipation and will not change the kinetic behavior of the model.

### II. COMPUTATIONAL AND ANALYTIC METHODS

#### A. Backward Master Equations and the Definitions of Speed and Accuracy

We quantify the speed and accuracy of the aminoacylation process using the framework of a first-passage process [20, 21]. Specifically, the temporal evolution of the system can be considered as Markovian jumps between discrete enzymatic states until the formation of a correct or incorrect product. In the tRNA^Ile^ aminoacylation network, we have 11 chemical states, which are labeled from 1 to 11 (see Figure S1). States 10 and 11 are end states which correspond to the formation of a correct and incorrect product, respectively. For each state *i*, we can define *F_R/W,i_*(*t*) as the first-passage probability density such that *F_R,i_*(*t*) d*t* (or *F_W,i_*(*t*) d*t*) quantifies the probability of forming a correct (or incorrect) product between *t* and *t* + d*t* without forming any products before time *t*, assuming that the enzyme starts at state *i* at time *t* = 0. In order quantify speed and accuracy, we define the splitting probability Π_*R/W*_ as the probability that the first product created is correct (incorrect), assuming that the system starts at state 1 (the free enzyme state E) at time *t* = 0:

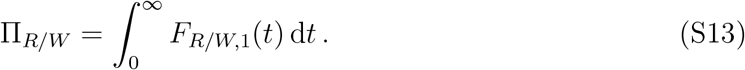

Naturally, accuracy is quantified by the error rate given by

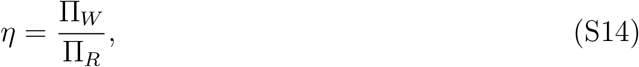

which is consistent with its traditional definition as the ratio between the product forming rates[22, 23]. The speed (i.e., inverse time) is quantified by the conditional mean first-passage time (MFPT), which is given by the (normalized) first moment of the first-passage probability density

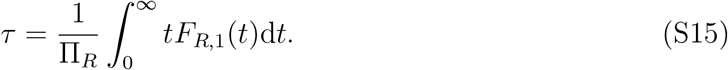

Now we present the mathematical formalism used to analytically determine *η* and *τ*. According to the definition, the first-passage probability densities of the end states read:

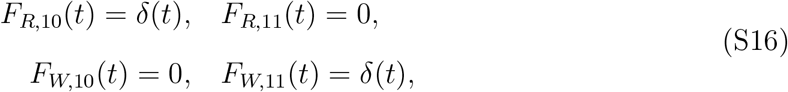

where *δ*(*t*) is the Dirac *δ* function. The time evolution of the first-passage probability densities of the other states *F_R/W,i_*(*t*) (*i* = 1, 2,…, 9) is governed by the backward master equations, which read:

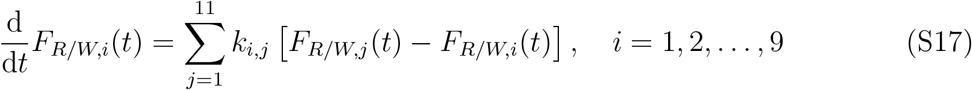

where *k_i,j_* denotes the first-order rate of transition from state *i* to state *j* and the initial conditions are *F_R/W,i_*(*t*) = 0 for *i* = 1, 2,…, 9. The transition rates *k_i,j_* in the tRNA^Ile^ aminoacylation network are compactly given by:

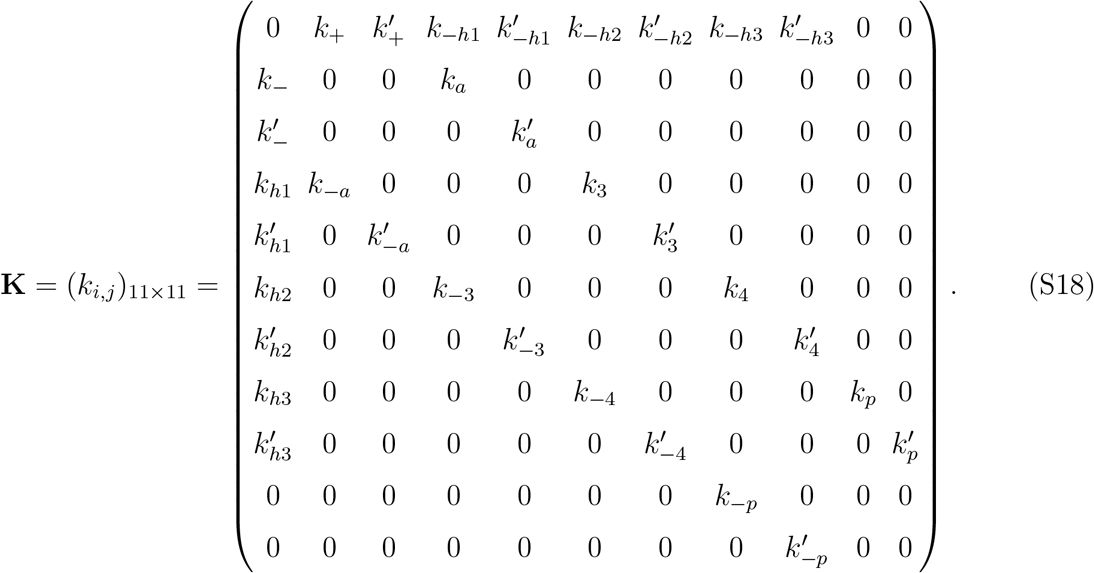

While Eqs. S16–S18 in principle can be directly solved for *F_R/W,i_*(*t*), it is usually more convenient to solve the backward master equations by performing a Laplace transformation 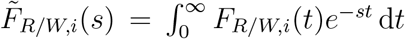. The transformed system obeys the following set of equations:

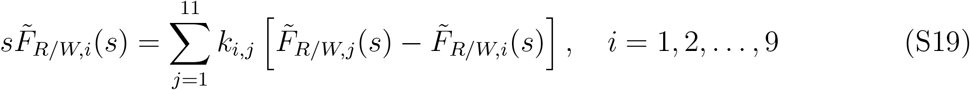

and

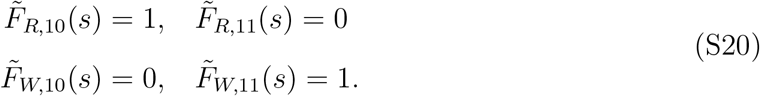

Equation S19 is a set of linear algebraic equations for 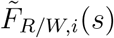 that can be solved analytically. The associated splitting probabilities Π_*R/W*_ are given by

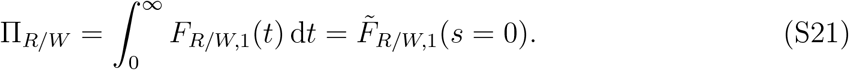

The error *η* is therefore determined as the ratio of the two splitting probabilities

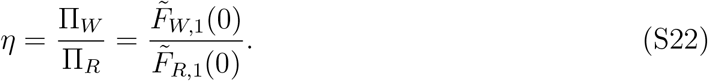

The conditional MFPT *τ* is given by the normalized first moment of the first-passage probability density:

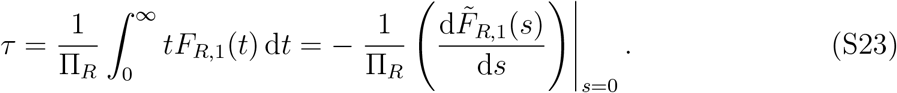

The symbolic linear algebra calculations based on Eqs. S17–S23 are done in Mathematica. The resulting expressions evaluated for the estimated parameter values are plotted in the main text figures.

#### B. Forward Master Equations and the Definition of Dissipation

In the forward framework, we use *P_i_*(*t*) to denote the probability of the enzyme being in chemical state *i* at time *t*. The main difference from the backward framework is that states 1, 10, and 11 should be considered to be the same state (i.e. the free enzyme). Therefore, we only have 9 distinct enzyme states. The probabilities of the enzyme being in different states are normalized by the following condition:

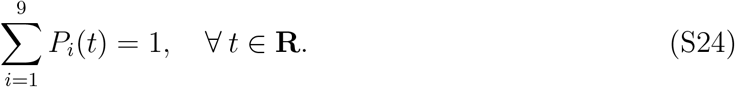

The time-dependent probability flux from state *i* to state *j* is given by *J_i,j_*(*t*) = *k_i,j_P_i_*(*t*). The time evolution of the probabilities *P_i_*(*t*) is governed by the forward master equations, which read:

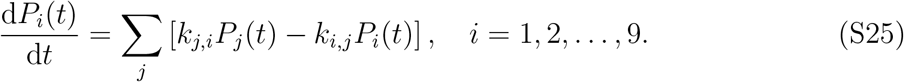

Note that merging states 1, 10, and 11 reduces the transition rates matrix **K** to a 9 × 9 matrix, which reads:

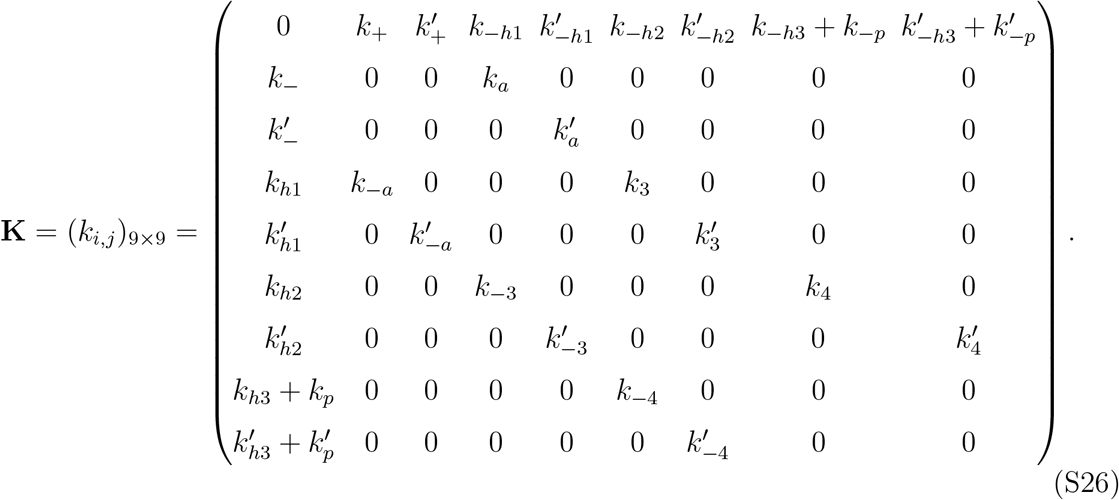

Setting the right hand side of Equation S25 to zero gives the steady-state probability distributions 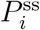. We can calculate the steady-state probability fluxes 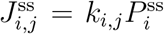 (The superscript *ss* will be dropped for simplicity). The steady state energy dissipation rate of the system is given by:

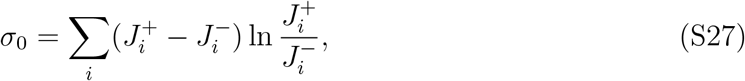

where 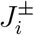 are the forward/backward steady-state fluxes of the chemical reaction *i*[19, 24].

It can be shown that the energy dissipation can be decomposed into three parts:

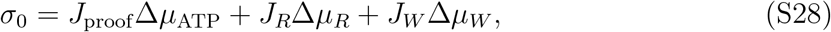

where *J*_proof_ is the sum of all net proofreading fluxes; *J_R_* and *J_W_* are the product formation fluxes for the right and wrong product, respectively. Δ*μ*_ATP_ = 29.5 *k_B_T* is the free energy released from ATP hydrolysis into AMP and PP_i_ under physiological concentrations of ATP, AMP, and PP_i_. Δ*μ_R_* = Δ*μ_p_* = 9.8 *k_B_T* is the free energy cost of forming a correctly charged isoleucyl-tRNA, namely the difference between the free energy released from the hydrolysis of ATP and that stored in the ester bond connecting tRNA and isoleucine. Δ*μ_W_* is the corresponding energy cost for valine, which is typically different from Δ*μ_R_*. However, since *J_W_* is much smaller than other fluxes, especially *J_R_*, the difference between Δ*μ_W_* and Δ*μ_R_* has negligible impact on the total dissipation. For the purpose of computing the total dissipation, we can safely ignore the last term in Eq. S28. We study the normalized dissipation rate *σ* defined as the dissipation rate per correct product formed:

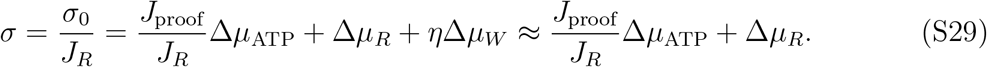

The symbolic linear algebra calculations based on Eqs. S24–S29 are done in Mathematica. The resulting expressions evaluated for estimated parameter values are plotted in the main text figures.

## Notes

### Competing Interest Statement

The authors have declared no competing interest.

### Summary of Updates

Added paragraph discussing the physical significance of the dissipation-error bound (Figure 4) in terms of proofreading fluxes for futile cycles in non-equilibrium systems.

